# A Biomechanical Hand Model to Quantify Finger Joint Kinematics Using a 3D Motion Capture System

**DOI:** 10.64898/2026.02.09.704796

**Authors:** Valeria Avilés-Carrillo, Ricardo G. Molinari, Guilherme A. G. De Villa, Leonardo A. Elias

**Affiliations:** Department of Electronics and Biomedical Engineering, School of Electrical and Computer Engineering, University of Campinas - UNICAMP, Campinas, 13083-852, SP, Brazil; Neural Engineering Research Laboratory, Center for Biomedical Engineering, University of Campinas - UNICAMP, Campinas, 13083-881, SP, Brazil; State University of Goiás - UEG, Itaberaí, 76630-000, GO, Brazil

**Keywords:** biomechanical model, finger movement kinematics, optical motion capture, metacarpophalangeal joint, thumb carpometacarpal joint

## Abstract

The kinematics of rhythmic, speed-modulated finger and grasp-like movements were analyzed using a reduced biomechanical model of the hand and a marker-based optical motion-capture system. Twenty-one healthy participants performed eight hand motor tasks involving metacarpophalangeal (MCP) joint flexion–extension (F–E) and carpometacarpal (CMC) thumb opposition–reposition (O–R) at two movement frequencies (0.50 and 0.75 Hz). Kinematic analysis quantified the range of movement (RoM), mean speed, and normalized total harmonic distortion (TDH_N_). Statistical analysis identified task type as the primary factor modulating all three metrics across digits, with large effect sizes 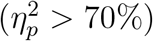. Movement frequency significantly influenced mean speed 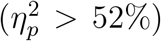 and moderately affected TDH_N_ 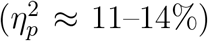, while thumb RoM remained statistically unchanged across frequencies (*p* = 0.063). Participants consistently reproduced the intended sinusoidal trajectories, as indicated by low TDH_N_ values (below 19%). The findings support the analysis of coordinated hand movements across various tasks under controlled time conditions. They also demonstrate that the simplified biomechanical model accurately captured both individual and co-ordinated finger movements. This provides a valuable reference for studies on motor control and for applications in rehabilitation and assistive technology.

## 1. Introduction

The human hand is one of the most complex and functionally versatile structures of the body, capable of performing movements that range from delicate manipulations to forceful grasps (Sobinov and Bensmaia, 2021). Its sophisticated anatomy, which integrates 27 bones, 29 joints, and over 120 ligaments, provides support and freedom of movement for the fingers. Additionally, a highly coordinated neural system integrates sensory and motor information to execute precise actions, depending on the task (Hirt et al., 2016; Bensmaia et al., 2020; Wang and Amirouche, 2025). The synergy between mechanical complexity and neuromuscular control has motivated extensive research on hand kinematics, aimed at describing finger movements in detail, quantifying their biomechanical parameters, and reproducing them through mathematical or robotic models (McFarland et al., 2023; Park et al., 2024; Ehara et al., 2025). Such efforts have generated key insights for rehabilitation (Bouteraa et al., 2023), prosthetic design (Lunguţ et al., 2023; Blana et al., 2020), and the development of human-machine interfaces (Zheng et al., 2021; García-Gil et al., 2025).

Previous studies have used optical and sensor-based motion capture systems to characterize finger kinematics in both healthy individuals and those with motor impairments, analyzing variables such as range of movement (RoM), peak angular velocity, joint sequencing, and temporal coordination during functional tasks including grasping, releasing, object transport, and other manipulative actions (Jarque-Bou et al., 2020; Xie et al., 2022; Li et al., 2023; Yuan et al., 2023; Gracia-Ibáñez et al., 2025). This body of work has shown that fingers do not contribute uniformly to hand kinematics: the thumb often exhibits distinct temporal patterns and sequencing, reflecting its specialized role in manual coordination (Gülke et al., 2012; Li et al., 2023), and both movement amplitude and velocity can vary substantially across tasks and conditions, even when the overall gesture appears similar (Kim et al., 2008; Bain et al., 2014; Klemm et al., 2024). Together, these findings highlight the complexity of inter-digit coordination and the task-dependent nature of motor control strategies.

However, most existing studies rely on kinematic descriptors that are primarily spatial and descriptive, and are typically obtained from static measurements or from dynamic tasks performed at self-selected speeds, without explicit control of movement rhythm. Consequently, they provide limited information about the temporal structure, spectral properties, and periodicity of rhythmic movements. In addition, there is a lack of systematic and comparative evaluations of multiple dynamic tasks performed under explicitly controlled temporal conditions, integrating individual finger movements and different grasping patterns within a single experimental protocol. This gap limits a unified understanding of how temporal constraints shape finger coordination and movement fidelity in rhythmic hand actions.

Therefore, the present study aimed to investigate periodic finger movements executed under explicit modulation of movement frequency imposed by a visual feedback interface in a healthy population. To this end, a reduced biomechanical model was employed, focusing on degrees of freedom (DoF) that are central to manual function, namely flexion–extension (F–E) at the metacarpophalangeal (MCP) joints of the index, middle, ring, and little fingers, and opposition–reposition (O–R) of the thumb at the carpometacarpal (CMC) joint. Kinematic data were collected while participants performed eight specific tasks at two different movement frequencies. This data was analyzed using commonly used kinematic variables, including range of motion (RoM), mean speed, and the normalized total harmonic distortion (THD_N_). The latter was used to assess how closely participants’ movements matched the sinusoidal visual command. By analyzing dynamic tasks in which participants reproduce prescribed trajectories in a controlled manner, this approach establishes kinematic references under well-defined temporal conditions. It facilitates comparisons across tasks, movement speeds, and levels of manual coordination, providing relevant information for studies of motor control, manual performance assessment, and rehabilitation applications.

## 2. Methods

### 2.1. Ethical Approval and Participants

Twenty-one volunteers (27.2 ± 6.5 years, 73.5 ± 16.1 kg, 1.68 ± 0.1 m) participated in the study after providing written informed consent. All participants self-reported as healthy with no prior history of neuromuscular diseases. The experimental protocol was conducted in accordance with the Declaration of Helsinki and approved by the Ethics Committee of the University of Campinas (CAAE 80299124.2.1001.5404) on 24 October 2024.

### 2.2. Experimental Protocol

The motor tasks used in the experimental protocol reproduced the four degrees of actuation of the Brunel 2.0 open-source bionic hand (Open Bionics, UK), encompassing both isolated finger movements and grasp-like actions. Task sequences were presented to participants via a virtual interface developed in Unity Engine (Unity Technologies, USA). The bionic hand was rendered as a 3D model, with joint rotations implemented using the Animator tool and Euler angles. This prosthetic-based visual interface was employed because the data collected for this study is part of a project aimed at designing a new hand prosthetic system.

The movement set included F–E of the index finger (Index), F–E of the middle finger (Middle), coupled F–E of the ring and little fingers (Ring-little), thumb O-R (Thumb), index–thumb pinch opening and closing (I.Pinch), middle–thumb pinch opening and closing (M.Pinch), tripod pinch involving the index, middle, and thumb fingers (Tripod), and five-finger hand opening and closing (5-Finger).

Each task lasted 45 seconds and was performed at two movement frequencies (0.50 Hz and 0.75 Hz), repeated across three sets, resulting in a total of 48 trials per participant. Movement sequences were randomized, with approximately 30 seconds of rest between repetitions. To ensure standardized execution, participants began each task with the hand fully opened and performed each pattern naturally. For thumb-related tasks, participants were instructed to completely reproduce the reposition action, as the thumb could otherwise remain in opposition while the other fingers moved. Fig. 1 illustrates the experimental setup and the performed tasks.

**Figure 1.**
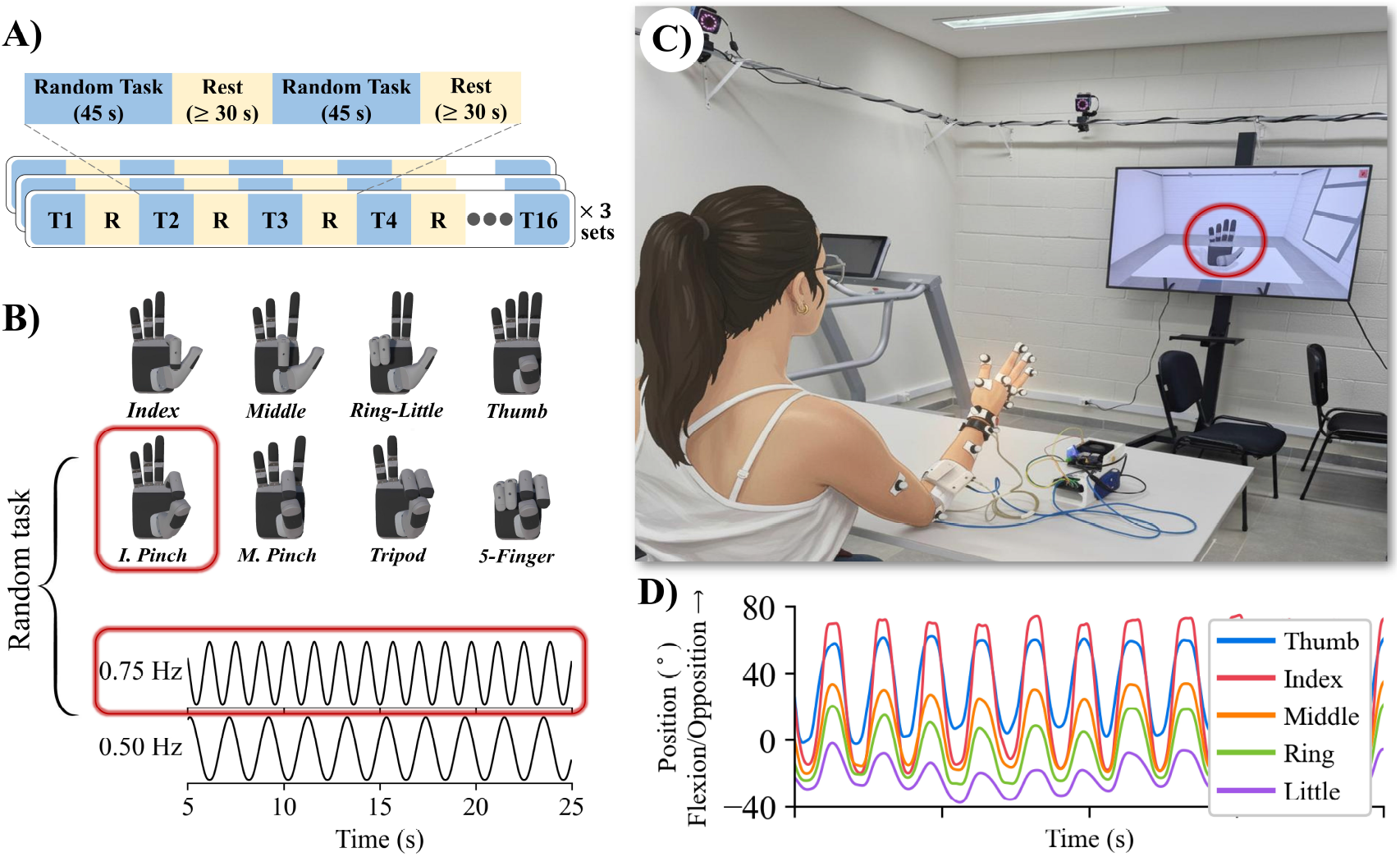
(A) Overview of the experimental protocol, comprising three sets of 16 randomized tasks interleaved with rest intervals. Draws randomized the order of tasks within each set without replacement from the combination of eight movement patterns and two movement frequencies. (B) Task patterns included flexion–extension (F-E) of the index, middle, and ring–little fingers; thumb opposition–reposition (O-R); and grasp-like movements (index pinch, middle pinch, tripod, and five-finger grip) performed at 0.75 Hz and 0.50 Hz. (C) Participants performed hand motor tasks while comfortably seated, with the elbow supported on a table, following on-screen sinusoidal finger movements of an animated bionic hand. (D) Finger joint kinematics for the I. Pinch task at 0.75 Hz, highlighted within the red boxes in (B).

### 2.3. Data Acquisition

Kinematic data were acquired using a Vicon Vero 2.2 motion capture system (Vicon Motion Systems, UK). The setup comprised eight high-speed infrared cameras operating at 100 frames per second, which recorded the positions of twelve 14 mm spherical retro-reflective markers on the arm, wrist, hand, and fingers (Fig. 2). The arrangement of wrist and hand reference points followed the upper-limb model established by Vicon Nexus (Vicon Motion Systems Limited, 2024), supplemented with nine additional points defined explicitly for the hand model. The anatomical locations for all markers are presented in Table 1.

**Table 1:** Anatomical landmarks for marker placement.

| Marker | Anatomical Positions |
| --- | --- |
| $P_T$ | At the head of the first metacarpal |
| $P_I$ | At the head of the second metacarpal |
| $P_M$ | At the head of the third metacarpal |
| $P_R$ | At the head of the fourth metacarpal |
| $P_L$ | At the head of the fifth metacarpal |
| $D_T^*$ | At the distal phalanx of the thumb |
| $D_I^*$ | At the distal phalanx of the index finger |
| $D_M^*$ | At the distal phalanx of the middle finger |
| $D_R^*$ | At the distal phalanx of the ring finger |
| $D_L^*$ | At the distal phalanx of the little finger |
| $L_W$ | At the styloid process of the ulna |
| $M_W$ | At the styloid process of the radius |

\* Markers placed on the nails of the corresponding fingers.

**Figure 2.**
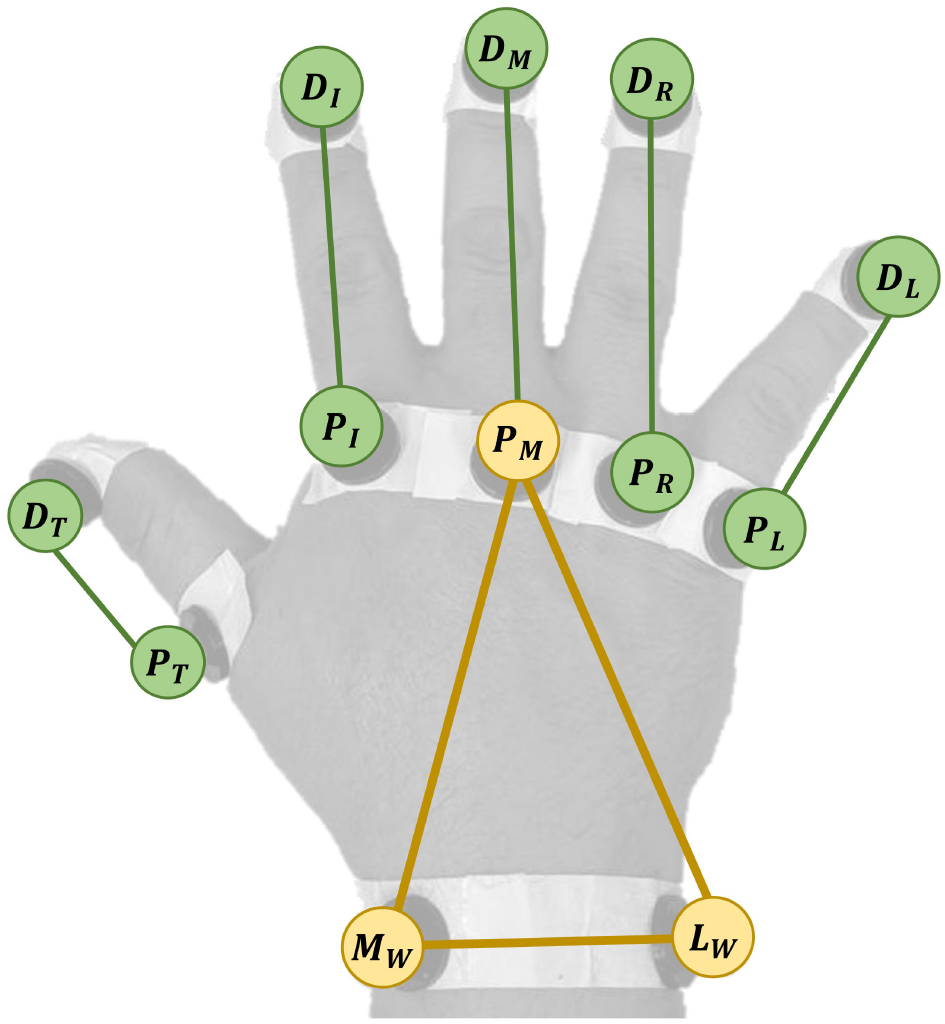
Arrangement of retro-reflective markers used in the Vicon motion capture system. Each finger received two markers: one on the distal phalanx (*D*_*f*_) and one on the metacarpal head (*P*_*f*_), where the subscript *f* denotes the first letter of the finger: index (*I*), middle (*M*), ring (*R*), little (*L*), and thumb (*T*). Wrist markers were placed on the medial (*M*_*W*_) and lateral (*L*_*W*_) sides of the joint.

Kinematic recordings were synchronized with the virtual bionic hand animation and high-density surface electromyography (HD sEMG) signals using a custom acquisition and processing system (Molinari et al., 2025). Synchronization triggers were transmitted wirelessly via TCP/IP to the animation interface and via UDP/IP to the motion capture system. Although the experimental protocol included HD sEMG acquisition, the analysis of these signals is out of the scope of the present work. The complete dataset, including both kinematic and HD sEMG data, is freely available (Avilés-Carrillo et al., 2026).

### 2.4. Marker Trajectory Reconstruction

Marker trajectories were reconstructed and labeled in 3D using Vicon Nexus 2.11.0 (Vicon Motion Systems, UK). Missing data were interpolated using cubic spline, cyclic, or pattern-based methods as appropriate, followed by a Woltring filter with a mean squared error of 5 mm^2^ (Woltring, 1986; Seegelke et al., 2011). Trials containing markers with gaps exceeding 200 frames (2 s) or in which the participant did not perform the task correctly were excluded from the analysis. A single trial was excluded from the entire dataset.

### 2.5. Biomechanical Hand Model

The biomechanical model was built using Vicon ProCalc version 1.6 (Vicon Motion Systems, UK) to calculate joint rotations and estimate F–E angles of the MCP joints for the index, middle, ring, and little fingers, as well as the O–R angle of the thumb CMC joint. The model uses the physical markers along with a virtual point (*C*_*W*_), defined as the midpoint between *M*_*W*_ and *L*_*W*_, to construct three vectors per finger 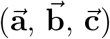.

For the index, middle, ring, and little fingers, 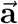connects the central wrist point (*C*_*W*_) to the metacarpal head (*P*_*f*_), while 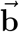 extends from the metacarpal head (*P*_*f*_) to the marker on the distal phalanx (*D*_*f*_). For the thumb, 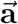 extends from the medial wrist point (*M*_*W*_) to the thumb metacarpal head (*P*_*T*_), and 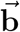 connects the lateral and medial wrist points (*L*_*W*_ → *M*_*W*_).

The 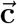 for the index, middle, and ring fingers was defined using the successive metacarpal heads: 2^nd^ → 3^rd^, 3^rd^ → 4^th^, and 4^th^ → 5^th^, respectively. For the little finger, 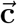 was defined from the 4^t h^ 5^t h^. For the thumb, 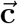 ex-tends from the medial wrist point to the index finger metacarpal head (*P*_*I*_). A detailed description of each vector is provided in Tab. 2.

**Table 2:** Definition of the vectors used in the hand model.

| Vector | Thumb | Index | Middle | Ring | Little |
| --- | --- | --- | --- | --- | --- |
| $\vec{a}$ | $M_W \rightarrow P_T$ | $C_W \rightarrow P_I$ | $C_W \rightarrow P_M$ | $C_W \rightarrow P_R$ | $C_W \rightarrow P_L$ |
| $\vec{b}$ | $L_W \rightarrow M_W$ | $P_I \rightarrow D_I$ | $P_M \rightarrow D_M$ | $P_R \rightarrow D_R$ | $P_L \rightarrow D_L$ |
| $\vec{c}$ | $M_W \rightarrow P_I$ | $P_M \rightarrow P_I$ | $P_R \rightarrow P_M$ | $P_L \rightarrow P_R$ | $P_L \rightarrow P_R$ |

Joint angles were computed as the rotation between vectors 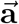 and 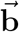 around the reference axis 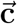. Both vectors were orthogonally projected onto the plane perpendicular to 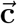 to remove components along the rotation axis (Eqs. (1)–(2)). The resulting projected vectors, 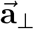 and 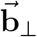, were normalized, and the angle between them was calculated using the dot product (Eq. (3)). Rotation direction was determined by the sign of 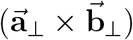 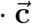, yielding an angle with the correct clockwise or counterclockwise orientation (Eq. (4)). The process ensures that the computed angle represents rotation exclusively about 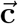 (Vicon Motion Systems Limited, 2022b,a). Fig. 3 illustrates the geometric definitions of the joint angles *θ*_*I*_ for the index finger and *θ*_*T*_ for the thumb.

**Figure 3.**
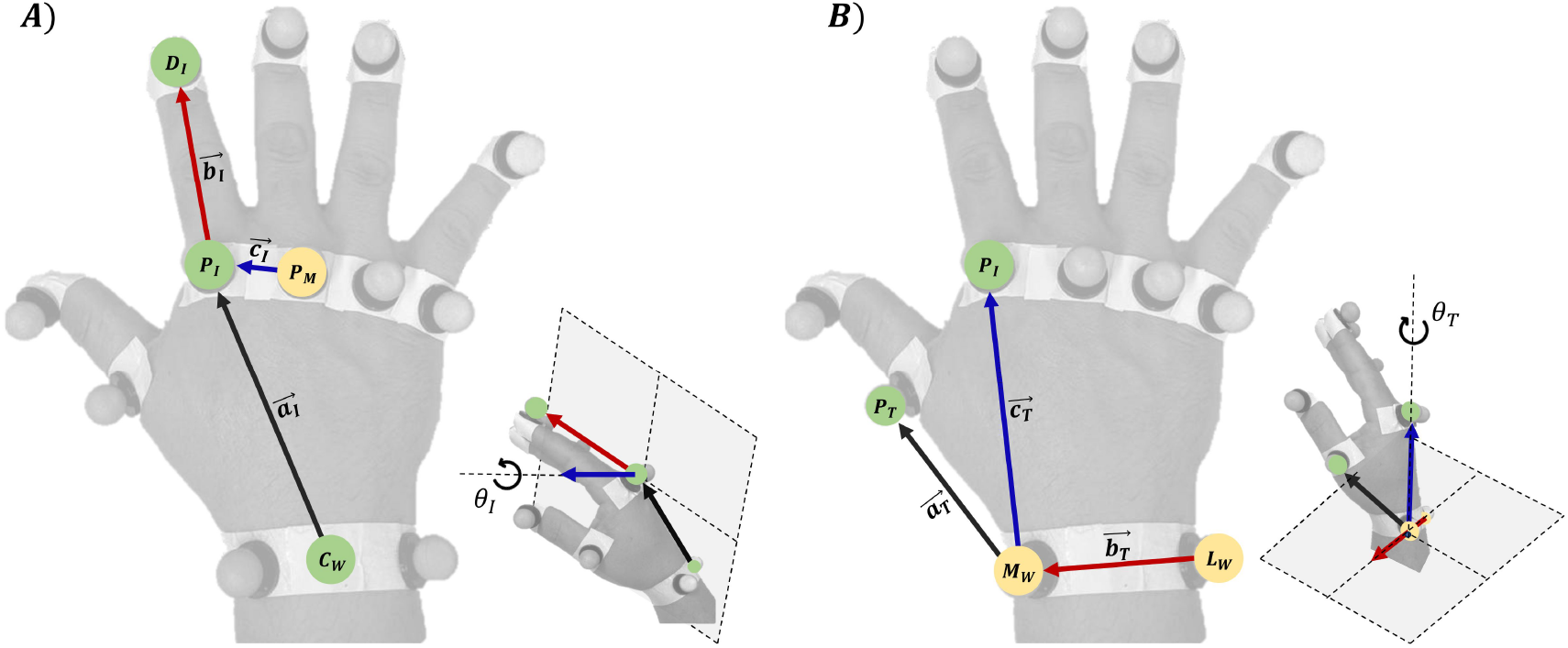
: Geometric representation of the vector configuration u sed to e stimate rotation angles at the MCP joint of the index finger (A) and the CMC joint of the thumb (B). Vectors are color-coded: 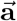 in black, 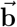 in red, and 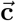 in blue. The projection plane, per-pendicular to 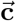, indicates the plane onto which vectors 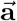 and 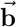 are projected to calculate the rotation angle.

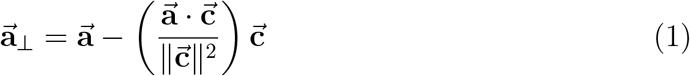

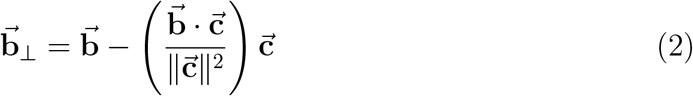

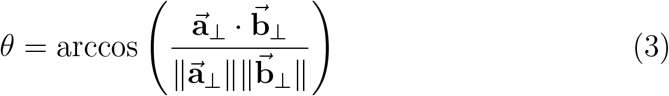

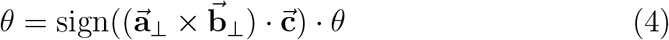

### 2.6. Data Analysis

The biomechanical model was evaluated using Python 3.12.7, applying three metrics to quantify differences across task types and frequencies and to assess movement accuracy. For each joint angle, the following parameters were calculated: (i) range of movement (RoM), measuring displacement amplitude; (ii) mean speed, representing movement dynamics; and (iii) normalized total harmonic distortion (TDH_N_) (Pietrosanti et al., 2023; Shimono et al., 2013), assessing adherence to the prescribed task and deviations from the expected pattern. Metric values were averaged across sets, providing a representative value per participant, finger, and task/frequency. The first and last five seconds of each recording were excluded to remove transient effects, and all trials were inspected for execution errors.

To calculate RoM, each task was divided into cycles by identifying discontinuities (wrap-around) in the Hilbert transform phase of the finger joint angle signal (Oppenheim and Schafer, 2009; Nayeem et al., 2021), which defined the start and end of each cycle. For multi-digit tasks, cycles were determined based on the finger with the highest interdependence (e.g., the ring finger for Ring-little, and the middle finger for Tripod) (Kim et al., 2008; Häger-Ross and Schieber, 2000a). For each cycle, the total number of samples and the starting sample were identified, enabling the segmentation of all five finger signals within the recording. Cycles containing fewer than 100 samples were discarded to avoid including partial cycles.

Subsequently, the RoM was defined as the average difference between the maximum and minimum angular positions within each cycle. Angular velocity was obtained by computing the discrete derivative of the angular position signal, calculated separately for each cycle. The mean speed was then estimated as the average of the mean absolute angular velocity across all cycles of each task.

The THD_N_, previously applied in its non-normalized form to periodic motor signals such as gait and pathological tremor (Martínez-Ramírez et al., 2015; Pietrosanti et al., 2023), was used here to assess deviations from an ideal periodic pattern across multiple DoFs and task conditions. To evaluate it, the power spectral density (PSD) of the angular position signals was estimated using the Welch method (Welch, 1967), with linear detrending and non-overlapping 8 s Hamming windows. The fundamental frequency (*f*_1_) was defined as the spectral component corresponding to the task frequency (0.50 Hz or 0.75 Hz), and the first four harmonics were subsequently identified. In addition, an analysis was conducted to verify whether the task was performed at the intended frequency by comparing the peak PSD component with *f*_1_. The amplitudes of the fundamental (*A*_1_) and the first four harmonics (*A*_2_–*A*_5_) were computed as described in Eq. (5), where *k* spans the three bins centered at the frequency of the *n*-th component (fundamental or harmonic), *Pxx*[*k*] denotes the PSD value in bin *k*, and Δ*f* is the spectral resolution (Retter et al., 2021; Cerna and Harvey, 2000).

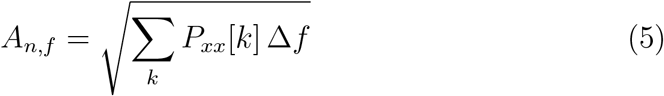

Eq. (6) defines the computation of TDH_N_ following the methods described in (Shmilovitz, 2005; Pietrosanti et al., 2025) and the normalization step described by Burden (2010). This normalization ensures that fingers with smaller ranges of motion yield TDH_N_ values proportional to their contribution, while more active fingers preserve values close to the original. The amplitude of the *n*-th harmonic of finger *f* is denoted as *A*_*n,f*_ (*n* = 2, 3, 4, 5). *A*_1,*f*_ indicates the amplitude of fundamental frequency for finger *f*, and *A*_1,max_ represents the maximum amplitude of fundamental frequency observed across all fingers in the trial.

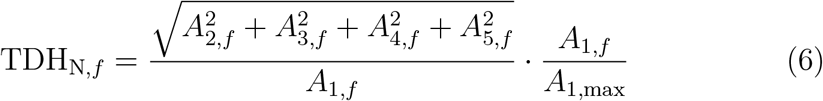

### 2.7. Statistical Analysis

Statistical analysis was conducted using the R programming language to examine differences in RoM, mean speed, and TDH_N_ across task types and frequencies for each finger. Normality was assessed with the Shapiro–Wilk test. Since the data violated this assumption, a two-way repeated-measures analysis of variance (Frequency × Task) was performed using the Aligned Rank Transform (ART) ANOVA procedure from the ARTool package. Posthoc pairwise comparisons of the estimated aligned rank means were adjusted using the Bonferroni correction, with significance set at *p <* 0.05. Effect sizes were calculated as partial eta squared 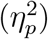 (Cohen, 1973) and interpreted as small 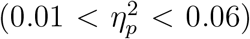, medium 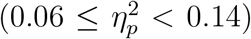, or large 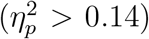(Mangiafico, 2016).

All results are reported as mean ± 95% confidence interval.

## 3. Results

Fig. 4 shows a representative example of the biomechanical model output during the Middle Pinch task at 0.75 Hz, along with the cycle-averaged joint angle, velocity, and position PSD (cycle duration ≈ 1.3 s). The estimated RoM were 60.4 ± 1.1 ^°^ (thumb), 43.3 ± 9.1 ^°^ (index), 91.1 ± 3.2 ^°^ (middle), 61.9 ± 3.6 ^°^ (ring), and 24.1 ± 4.4 ^°^ (little finger), with corresponding mean angular speeds of 69.8 ± 1.9 ^°^/s, 48.6 ± 9.8 ^°^/s, 106.1 ± 6.0 ^°^/s, 71.3 ± 3.8 ^°^/s, and 27.8 ± 4.9 ^°^/s. The thumb and middle fingers, which were directly involved in the task, exhibited consistent trajectories with low cycle-to-cycle variance. In contrast, the index, ring, and little fingers showed greater variability, with the ring displaying intermediate variance, likely reflecting its mechanical coupling to the middle finger. The thumb and middle fingers reached the highest speeds, followed by the ring finger, whereas the index and little fingers showed lower values.

**Figure 4.**
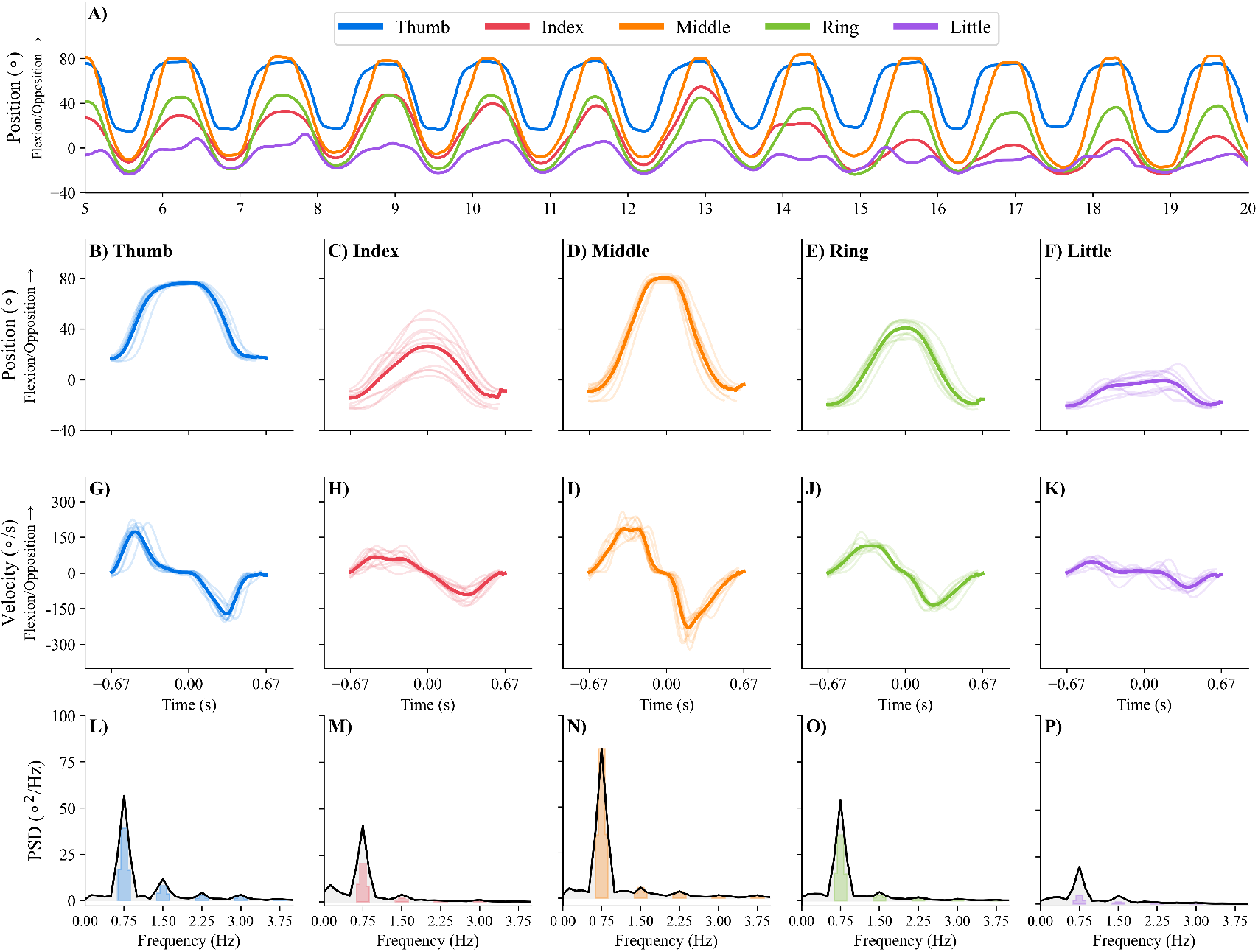
: Illustrative example of kinematic data and power spectral density (PSD) from a participant performing the Middle Pinch task at 0.75 Hz. (A) Angular positions of all five fingers over a 20-s segment extracted from the 45-s recording, where increasing values indicate finger flexion and thumb opposition. (B–F) Cycle-averaged position trajectories for the thumb to little finger. (G–K) Instantaneous movement velocities. Thin, transparent lines represent individual cycles, and their averages are shown as thick, solid lines. (L–P) PSD for the thumb to the little fingers. Shaded areas indicate the frequency bins used to estimate the fundamental frequency (*f*_1_) and its harmonics. Bar height reflects normalized amplitudes.

The fundamental component (*f*_1_) matched the task frequency (0.75 Hz) across all measured movements. TDH_N_ values were 16 %, 6 %, 10 %, 7 %, and 6 % for the thumb to little fingers, respectively. The results indicate that only 6 % to 16 % of the total signal power was contained in the harmonics. At the same time, most of the power was concentrated in the fundamental component (*f*_1_), suggesting that the participant closely followed the visual command (Pietrosanti et al., 2025).

### 3.1. Range of Movement

The ART ANOVA on RoM revealed that task was the primary factor influencing movement amplitude across all digits (*p <* 0.001, 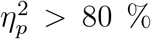). Task frequency also influenced RoM, with higher values at 0.75 Hz compared to 0.50 Hz. The effect was significant but small for the middle and little fingers (*p <* 0.001, 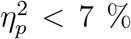) and medium-sized for the index (*p <* 0.001, 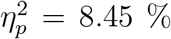) and ring fingers (*p <* 0.001, 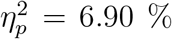). In contrast, the thumb did not exhibit statistically significant differences across frequencies (*p* = 0.063). No significant interactions between frequency and task were observed for RoM. Detailed statistical results are presented in Tab. 3.

**Table 3:** *p*-values and effect sizes 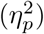for range of movement (RoM), mean speed, and normalized total harmonic distortion (TDH_N_) for each finger.

| Finger | Factor | RoM |  | Mean speed |  | TDH<sub>N</sub> |  |
| --- | --- | --- | --- | --- | --- | --- | --- |
| | | $p$-value | $\eta_p^2$ (%) | $p$-value | $\eta_p^2$ (%) | $p$-value | $\eta_p^2$ (%) |
| Thumb | Freq. | 0.063 | 1.1 | < **0.001** | 52.0 | < **0.001** | 12.1 |
|  | Task | < **0.001** | 85.1 | < **0.001** | 85.0 | < **0.001** | 83.1 |
| | Freq. × Task | 0.462 | 2.1 | < **0.001** | 36.1 | **0.001** | 7.3 |
| Index | Freq. | < **0.001** | 8.4 | < **0.001** | 76.5 | < **0.001** | 13.2 |
|  | Task | < **0.001** | 94.5 | < **0.001** | 94.4 | < **0.001** | 86.1 |
| | Freq. × Task | 0.568 | 1.8 | < **0.001** | 50.3 | **0.048** | 4.5 |
| Middle | Freq. | < **0.001** | 5.8 | < **0.001** | 65.5 | < **0.001** | 13.0 |
|  | Task | < **0.001** | 85.2 | < **0.001** | 85.3 | < **0.001** | 84.7 |
| | Freq. × Task | 0.557 | 1.9 | < **0.001** | 32.7 | 0.147 | 3.5 |
| Ring | Freq. | < **0.001** | 6.9 | < **0.001** | 63.4 | < **0.001** | 13.8 |
|  | Task | < **0.001** | 91.5 | < **0.001** | 91.8 | < **0.001** | 83.6 |
| | Freq. × Task | 0.164 | 3.3 | < **0.001** | 39.5 | **0.033** | 4.9 |
| Little | Freq. | < **0.001** | 5.4 | < **0.001** | 55.3 | < **0.001** | 11.0 |
|  | Task | < **0.001** | 83.6 | < **0.001** | 83.0 | < **0.001** | 76.9 |
| | Freq. × Task | 0.405 | 2.3 | < **0.001** | 43.2 | 0.097 | 3.9 |

Results for the main factors (frequency and task) and their interaction are shown.
Bold values indicate statistical significance at $p < 0.05$ .

Post-hoc pairwise comparisons showed that fingers exhibited greater RoM during tasks in which they were actively engaged compared to conditions in which their movements were not intentional, producing distinct patterns of coordinated RoM with significant differences in most comparisons. When a finger was directly involved, differences decreased in most cases, except for the ring, which retained statistical significance across all conditions. Additionally, except for the thumb, non-involved adjacent digits showed progressively reduced RoM with increasing distance from the primary active digits. All pairwise comparisons are provided in the Appendix A.

Particularly, the thumb RoM ranged from 45^°^ to 59^°^ when actively engaged and remained near 5^°^–7^°^ when inactive, reaching its highest values during the O-R task (Thumb) and pinch movements (I. Pinch, M. Pinch). The index and middle fingers attained maximum RoM during individual F-E tasks, whereas the ring and little fingers peaked during the simultaneous F-E task (Ring-little). Descriptive statistics of RoM across all participants, grouped by finger, task type, and task frequency, are summarized in Tab. 4 and Fig. 5 (A-E).

**Table 4:** Descriptive data (mean ± 95% confidence interval) for range of movement (RoM), mean speed, and normalized total harmonic distortion (TDH_N_) for each task type, task frequency (0.50 Hz and 0.75 Hz), and finger.

| Task | RoM (°) |  |  |  |  |  |  |  |  |  |
| --- | --- | --- | --- | --- | --- | --- | --- | --- | --- | --- |
|  | Thumb |  | Index |  | Middle |  | Ring |  | Little |  |
|  | 0.50 Hz | 0.75 Hz | 0.50 Hz | 0.75 Hz | 0.50 Hz | 0.75 Hz | 0.50 Hz | 0.75 Hz | 0.50 Hz | 0.75 Hz |
| Index | 7.61 ± 2.26 | 7.04 ± 2.59 | 114.67 ± 6.97 | 122.00 ± 5.75 | 44.87 ± 8.68 | 48.68 ± 8.65 | 28.37 ± 7.14 | 30.04 ± 6.99 | 13.77 ± 4.56 | 14.75 ± 4.53 |
| Middle | 5.33 ± 2.28 | 5.73 ± 2.33 | 43.61 ± 7.41 | 48.13 ± 6.57 | 111.30 ± 8.85 | 117.04 ± 7.39 | 64.96 ± 7.60 | 72.19 ± 8.46 | 21.70 ± 4.74 | 24.05 ± 5.03 |
| Ring-little | 5.29 ± 2.32 | 5.85 ± 2.42 | 13.96 ± 3.58 | 17.25 ± 3.92 | 50.27 ± 10.12 | 59.41 ± 10.63 | 97.93 ± 7.75 | 109.11 ± 8.28 | 97.04 ± 8.14 | 108.44 ± 8.72 |
| Thumb | 55.19 ± 6.21 | 59.47 ± 5.96 | 5.71 ± 1.24 | 7.03 ± 2.16 | 3.95 ± 1.07 | 4.72 ± 1.74 | 3.92 ± 0.79 | 4.50 ± 1.17 | 4.75 ± 1.11 | 5.45 ± 1.76 |
| I. Pinch | 45.39 ± 6.20 | 46.83 ± 6.01 | 77.41 ± 6.38 | 83.78 ± 6.41 | 32.49 ± 6.54 | 37.75 ± 6.56 | 19.37 ± 3.89 | 22.98 ± 4.09 | 11.63 ± 2.47 | 13.78 ± 2.87 |
| M. Pinch | 48.31 ± 5.96 | 49.50 ± 6.55 | 37.18 ± 5.17 | 42.46 ± 4.52 | 72.63 ± 5.28 | 77.45 ± 4.89 | 48.41 ± 5.00 | 52.62 ± 5.17 | 19.04 ± 3.24 | 22.16 ± 3.47 |
| Tripod | 49.95 ± 6.14 | 50.52 ± 6.49 | 79.37 ± 5.73 | 83.50 ± 4.78 | 78.13 ± 5.66 | 82.42 ± 4.84 | 54.13 ± 5.74 | 58.43 ± 5.19 | 21.66 ± 4.08 | 23.82 ± 4.28 |
| 5-Finger | 53.94 ± 6.45 | 54.29 ± 6.83 | 78.79 ± 5.12 | 84.38 ± 4.79 | 72.62 ± 5.07 | 78.26 ± 4.65 | 70.82 ± 4.91 | 76.06 ± 4.59 | 72.22 ± 5.62 | 77.07 ± 4.83 |
| Task | Mean Speed (°/s) |  |  |  |  |  |  |  |  |  |
|  | Thumb |  | Index |  | Middle |  | Ring |  | Little |  |
|  | 0.50 Hz | 0.75 Hz | 0.50 Hz | 0.75 Hz | 0.50 Hz | 0.75 Hz | 0.50 Hz | 0.75 Hz | 0.50 Hz | 0.75 Hz |
| Index | 7.87 ± 2.26 | 10.31 ± 3.75 | 117.54 ± 6.52 | 183.90 ± 8.72 | 45.27 ± 8.58 | 72.13 ± 13.18 | 28.55 ± 7.11 | 44.28 ± 10.55 | 13.60 ± 4.54 | 21.25 ± 6.73 |
| Middle | 5.74 ± 2.23 | 8.66 ± 3.41 | 44.97 ± 7.12 | 72.11 ± 9.83 | 115.96 ± 7.40 | 177.11 ± 10.24 | 67.30 ± 7.17 | 108.72 ± 12.31 | 22.25 ± 4.79 | 35.77 ± 7.60 |
| Ring-little | 5.56 ± 2.35 | 8.58 ± 3.54 | 14.85 ± 3.72 | 25.59 ± 5.73 | 53.75 ± 10.44 | 89.69 ± 15.77 | 104.81 ± 7.51 | 166.05 ± 11.82 | 103.71 ± 8.05 | 164.53 ± 12.41 |
| Thumb | 60.41 ± 5.69 | 91.60 ± 9.32 | 6.42 ± 1.38 | 10.61 ± 3.22 | 4.12 ± 1.11 | 6.96 ± 2.62 | 4.16 ± 0.81 | 6.66 ± 1.74 | 4.96 ± 1.16 | 8.04 ± 2.72 |
| I. Pinch | 46.77 ± 6.39 | 69.98 ± 9.24 | 78.98 ± 5.89 | 125.06 ± 9.35 | 32.51 ± 6.43 | 55.43 ± 9.82 | 19.26 ± 3.73 | 33.47 ± 6.05 | 11.37 ± 2.29 | 19.60 ± 4.14 |
| M. Pinch | 50.38 ± 6.28 | 74.27 ± 9.86 | 37.86 ± 5.10 | 63.42 ± 6.67 | 74.33 ± 4.76 | 116.09 ± 7.22 | 49.32 ± 4.78 | 78.63 ± 7.67 | 19.16 ± 3.30 | 32.53 ± 5.13 |
| Tripod | 51.00 ± 6.24 | 74.81 ± 9.75 | 80.81 ± 5.57 | 124.64 ± 6.99 | 79.34 ± 5.50 | 122.81 ± 7.08 | 54.59 ± 5.62 | 86.63 ± 7.60 | 21.54 ± 4.11 | 34.59 ± 6.33 |
| 5-Finger | 54.97 ± 6.63 | 80.98 ± 10.20 | 79.62 ± 4.91 | 126.36 ± 7.11 | 73.28 ± 4.84 | 117.19 ± 6.93 | 71.29 ± 4.62 | 113.70 ± 6.71 | 72.42 ± 5.35 | 114.65 ± 7.10 |
| Task | TDH<sub>N</sub> (%) |  |  |  |  |  |  |  |  |  |
|  | Thumb |  | Index |  | Middle |  | Ring |  | Little |  |
|  | 0.50 Hz | 0.75 Hz | 0.50 Hz | 0.75 Hz | 0.50 Hz | 0.75 Hz | 0.50 Hz | 0.75 Hz | 0.50 Hz | 0.75 Hz |
| Index | 1.85 ± 0.51 | 1.22 ± 0.41 | 16.38 ± 2.54 | 14.30 ± 1.17 | 6.84 ± 1.19 | 5.30 ± 0.85 | 5.06 ± 1.18 | 3.84 ± 0.99 | 2.65 ± 0.98 | 2.00 ± 0.86 |
| Middle | 1.44 ± 0.36 | 1.04 ± 0.27 | 6.70 ± 1.29 | 5.51 ± 1.06 | 15.51 ± 2.04 | 13.47 ± 1.70 | 9.61 ± 1.65 | 8.43 ± 1.33 | 3.89 ± 0.63 | 3.27 ± 0.61 |
| Ring-little | 1.25 ± 0.34 | 0.93 ± 0.27 | 2.92 ± 0.66 | 2.12 ± 0.49 | 8.36 ± 1.69 | 6.63 ± 1.15 | 16.13 ± 2.26 | 12.39 ± 1.49 | 16.49 ± 2.42 | 12.72 ± 1.50 |
| Thumb | 19.39 ± 2.72 | 17.54 ± 2.70 | 4.01 ± 1.23 | 3.59 ± 1.35 | 2.18 ± 0.77 | 2.01 ± 0.91 | 2.15 ± 0.58 | 1.59 ± 0.43 | 2.63 ± 0.80 | 2.17 ± 0.91 |
| I. Pinch | 12.58 ± 2.77 | 10.51 ± 2.73 | 16.57 ± 2.31 | 14.57 ± 2.49 | 6.77 ± 1.59 | 5.85 ± 1.69 | 4.37 ± 0.89 | 3.87 ± 0.94 | 3.09 ± 0.70 | 2.66 ± 0.75 |
| M. Pinch | 15.97 ± 2.96 | 12.85 ± 2.59 | 8.46 ± 1.32 | 7.23 ± 1.53 | 15.56 ± 2.25 | 13.83 ± 2.44 | 10.71 ± 1.57 | 9.12 ± 1.67 | 5.38 ± 1.10 | 4.34 ± 0.86 |
| Tripod | 14.81 ± 2.82 | 12.42 ± 2.64 | 16.00 ± 1.81 | 13.78 ± 2.35 | 15.72 ± 1.79 | 13.65 ± 2.37 | 11.28 ± 1.57 | 9.44 ± 1.59 | 5.47 ± 1.00 | 4.42 ± 0.74 |
| 5-Finger | 17.47 ± 3.16 | 14.61 ± 2.94 | 16.52 ± 2.20 | 13.48 ± 1.88 | 15.39 ± 2.30 | 12.75 ± 1.92 | 15.52 ± 2.10 | 12.81 ± 1.63 | 15.63 ± 2.18 | 12.78 ± 1.78 |

**Figure 5.**
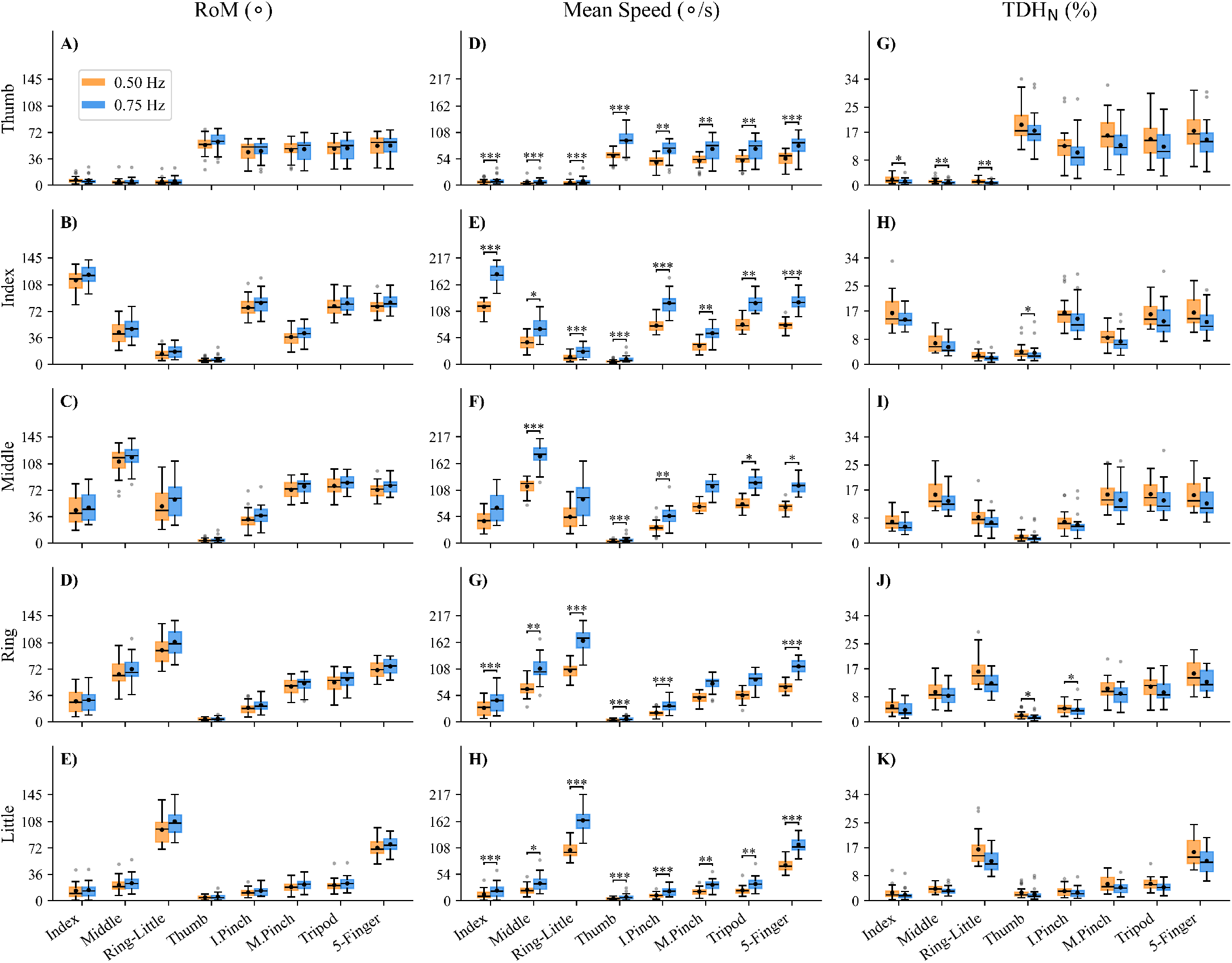
: Distribution of range of movement (RoM), mean speed, and normalized total harmonic distortion (TDH_N_) grouped by task type, task frequency, and finger (*N* = 62). Panels (A–E), (F–H), and (I–K) display RoM, mean speed, and TDH_N_ for the thumb to little finger, respectively. Asterisks indicate post-hoc pairwise comparison differences between frequency levels within each task following a significant Task × Frequency interaction: *p <* 0.05 (∗), *p <* 0.01 (∗∗), and *p <* 0.001 (∗∗∗). Black dots represent mean values, boxes indicate the interquartile range, whiskers show minimum and maximum values (excluding outliers), and gray circles denote outliers.

### 3.2. Mean Speed

The ART ANOVA on mean speed revealed significant effects of task, frequency, and their interaction, all with large effect sizes (*p <* 0.001, 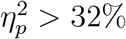; Tab. 3). Post-hoc pairwise comparisons were performed to assess differences in frequency within each task, as the RoM analysis indicated a strong effect of task type on finger motion. Overall, mean speed increased from 0.50 Hz to 0.75 Hz, although the magnitude of this effect varied depending on the executed gesture and the specific digit.

The thumb, index, and little fingers showed significant increases in mean speed with task frequency across all tasks (Fig. 5D-H). The middle finger also exhibited clear increases in most cases, except for Index (*p* = 0.055), M. Pinch (*p* = 0.053), and Ring–Little (*p* = 0.760) tasks, where differences did not reach statistical significance. For the ring finger, no significant changes were observed in the M. Pinch (*p* = 0.906) and Tripod (*p* = 0.448). Complete results of the post-hoc pairwise comparisons for mean speed are provided in the Appendix B.

### 3.3. Normalized Total Harmonic Distortion

The ART ANOVA on TDH_N_ revealed significant differences (*p <* 0.001) between tasks (large effect size, 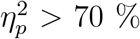) and frequencies (medium effect size, 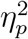 between 11 % and 14 %) across all digits, as shown in Tab. 3. Frequency × task interactions were significant only for the thumb, index, and ring fingers (*p <* 0.001), with small to medium effect sizes 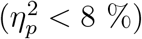.

TDH_N_ remained low across all conditions (Fig. 5G–K), with the smallest values observed in fingers not involved in the task. Overall, significant differences (*p <* 0.05) were observed between conditions involving active versus uninvolved fingers, across both task and frequency levels. Regarding the interaction effects, TDH_N_ for the thumb was higher at 0 .50 Hz than at 0.75 Hz in the Index, Middle, and Ring–Little tasks, with mean values below 2 %. The index finger showed a significant difference only in the Thumb task (mean values below 5 %), whereas the ring finger exhibited changes in the Thumb and I. Pinch tasks. Complete results of all pairwise comparisons are provided in the Appendix C.

Even for fingers actively involved in the tasks, the normalized TDH_N_ remained low under all conditions, below 19% for the thumb, 16% for the index, 15% for the middle, 16% for the ring, and 16% for the little finger, indicating consistent task performance.

Across all recordings, eleven trials presented peak PSD values that did not coincide with the target task frequency, suggesting minor deviations in movement execution (Tab. 5). In two cases (Index 0.50 Hz and Thumb 0.75 Hz), the participant did not start with the hand fully open. In six others (Index 0.50 Hz, Thumb 0.50 Hz, Tripod 0.50 Hz, Index 0.75 Hz, Thumb 0.75 Hz, and Index Pinch 0.75 Hz), participants skipped movement cycles, performed cycles with reduced amplitude, or executed them at a higher speed. The remaining cases, in which the detected peak PSD deviated from the target frequency, showed no evident execution issues and were likely due only to being out of synchronization with the virtual bionic hand movement.

**Table 5:** Mean ± 95% confidence interval (*N*) of the fundamental frequency extracted from the PSD, computed only for trials in which the detected frequency did not match the target task frequency.

| 0.50 Hz task frequency |  |  |  |  |  |  |  |  |  |  |
| --- | --- | --- | --- | --- | --- | --- | --- | --- | --- | --- |
| Task | Thumb |  | Index |  | Middle |  | Ring |  | Little |  |
| | Value | $N$ | Value | $N$ | Value | $N$ | Value | $N$ | Value | $N$ |
| Index | $0.23 \pm 0.08$ | (15) | **0.75** | (1) | 0.75 | (1) | 0.75 | (1) | $0.31 \pm 0.23$ | (6) |
| Middle | $0.19 \pm 0.04$ | (26) | $0.62 \pm 0.00$ | (2) | **$0.62 \pm 0.00$** | (2) | $0.62 \pm 0.00$ | (2) | $0.62 \pm 0.00$ | (2) |
| Ring-little | $0.25 \pm 0.09$ | (25) | $0.28 \pm 0.18$ | (5) | 0.62 | (1) | — | | — | |
| Thumb | **0.62** | (1) | $0.27 \pm 0.09$ | (33) | $0.23 \pm 0.08$ | (35) | $0.18 \pm 0.05$ | (25) | $0.20 \pm 0.05$ | (27) |
| I. Pinch | — | | — | | — | | 0.12 | (1) | $0.12 \pm 0.00$ | (3) |
| M. Pinch | — |  | — |  | — |  | — |  | — |  |
| Tripod | **0.62** | (1) | **0.62** | (1) | **0.62** | (1) | 0.62 | (1) | $0.38 \pm 0.49$ | (2) |
| 5-Finger | **0.62** | (1) | **0.62** | (1) | **0.62** | (1) | **0.62** | (1) | **0.62** | (1) |
| 0.75 Hz task frequency |  |  |  |  |  |  |  |  |  |  |
| Task | Thumb |  | Index |  | Middle |  | Ring |  | Little |  |
| | Value | $N$ | Value | $N$ | Value | $N$ | Value | $N$ | Value | $N$ |
| Index | $0.34 \pm 0.20$ | (20) | **1.00** | (1) | 1.00 | (1) | 1.00 | (1) | $0.25 \pm 0.17$ | (10) |
| Middle | $0.21 \pm 0.08$ | (19) | — | | — | | — | | — | |
| Ring-little | $0.17 \pm 0.05$ | (20) | $0.50 \pm 0.73$ | (2) | 0.88 | (1) | **0.88** | (1) | **0.88** | (1) |
| Thumb | **$0.81 \pm 0.37$** | (2) | $0.35 \pm 0.15$ | (34) | $0.29 \pm 0.15$ | (31) | $0.17 \pm 0.04$ | (25) | $0.17 \pm 0.06$ | (23) |
| I. Pinch | **0.88** | (1) | **0.88** | (1) | 0.88 | (1) | 0.88 | (1) | $0.31 \pm 0.37$ | (4) |
| M. Pinch | — |  | — |  | — |  | — |  | 0.12 | (1) |
| Tripod | — |  | — |  | — |  | — |  | — |  |
| 5-Finger | — |  | — |  | — |  | — |  | — |  |

Bold values indicate fingers actively engaged in the task.
Dashes (—) indicate that all trials matched the expected frequency for that task-finger combination.

## 4. Discussion

The proposed biomechanical model quantified finger-joint kinematics, focusing on thumb O-R at the CMC joint and F-E at the MCP joints. Participants performed eight tasks involving individual and combined finger movements under controlled temporal conditions at two speeds. A detailed characterization of movement patterns across tasks and fingers was obtained by analyzing kinematic features including RoM, mean speed, and TDH_N_. The results show reproducible and distinct kinematic patterns across tasks, highlight the reduced biomechanical model’s ability to capture finger movements, and provide a valuable reference for motor control research and the development of human–machine interfaces.

### 4.1. Range of Movement and Speed

The MCP F–E RoM values of the index to little fingers during cyclostationary movements at 0.50 and 0.75 Hz were broadly consistent with prior reports, despite methodological differences. Relative to the large-scale goniometric study by (B.K et al., 2024), slightly higher mean values were observed for the index and middle fingers, whereas the ring and little fingers showed comparable or smaller ranges. Consistently, (Reissner et al., 2019) reported MCP RoM values of the same order using 3D motion capture, which demonstrated higher test–retest reliability and lower measurement error than manual goniometry. In addition, (Li et al., 2023) found mean MCP RoM amplitudes between 72^°^ and 77^°^ for the same four fingers during maximal-speed dynamic opening–closing 5-finger task, closely matching values observed in our study for this grasping movement at both frequencies. Overall, the observed agreement in RoM values, together with the lower computational complexity of the proposed approach, supports the validity of MCP F–E RoM measurements during cyclostationary tasks and highlights the advantages of motion-capture-based assessment for functional kinematic mapping and algorithm training in bionic systems (He et al., 2019; Moeinnia et al., 2022).

For the thumb, active O–R RoM values were consistent with previously reported data for the CMC joint in healthy subjects (≈61^°^ for anteposition and radial abduction; (Barakat et al., 2013)). The study demonstrated high accuracy and reproducibility for most thumb measurements, while revealing that reductions in specific joint movements, such as MCP extension, may be compensated by increased motion in other joints. Given that thumb opposition integrates abduction, flexion, and medial rotation, the present model provides an approximate estimation of CMC motion, capturing the inherent complexity of thumb kinematics and the compensatory interplay between joints. Importantly, measurements based on skin markers or goniometry primarily reflect overall thumb motion and may over- or underestimate true CMC rotations, underscoring the value of this approach as an informative assessment of CMC kinematics (Crisco et al., 2025). Moreover, the thumb exhibited greater sensitivity to task demands relative to high-speed dynamic tasks (Li et al., 2023), with ≈54^°^ measured in our study compared with ≈26^°^ reported during rapid movements. The difference likely reflects variations in task execution, as Li et al. performed rapid maximal-speed cycles while our study employed cyclostationary movements at controlled frequencies. Overall, the findings suggest that MCP joint kinematics are relatively consistent across task conditions, whereas thumb RoM is more sensitive to task-specific constraints and movement strategy.

Having established the validity of our measurements, we next examined the influence of movement speed on joint RoM. The increase in RoM at faster movement speeds for the index, middle, ring, and little fingers likely arises from the dynamic properties of the musculoskeletal system rather than from primary neural adaptation. Higher movement velocities amplify segment momentum, and motion termination increasingly relies on passive tissue tension. Similar patterns have been observed in externally paced tasks (Häger-Ross and Schieber, 2000b), where faster movements produced larger joint excursions and a more even distribution across the finger kinematic chain. Consequently, higher kinetic energy combined with shorter movement duration results in larger joint displacements (Orendurff et al., 2018; Yang et al., 2025).

### 4.2. Normalized Total Harmonic Distortion

Participants closely replicated the sinusoidal pattern of the visual feed-back from the virtual bionic prosthetic hand, with low harmonic distortion. Normalizing total harmonic distortion facilitated interpretation, as greater TDH_N_ magnitudes reflect deviations weighted by each finger’s relative contribution to the total movement amplitude. Notably, the metric decreased at 0.75 Hz compared to 0.50 Hz, despite the observed increase in RoM and mean speed at higher frequencies. The identified pattern likely reflects the combined effects of passive, inertial, and elastic properties of the musculoskeletal system, which act as a natural biomechanical filter (Salmond et al., 2017; Binder-Markey and Murray, 2017). The reduced time available for corrective adjustments at faster movement rates further limits deviations, thereby smoothing angular trajectories (Denyer and Boyd, 2025; Guérin et al., 2021).

### 4.3. Future Work

Future investigations will build upon the present biomechanical model and the data already obtained to advance further its application in the training and validation of control algorithms for prosthetic and robotic devices. In this context, future developments will incorporate thumb flexion and extension, as well as wrist movements, which are predominantly driven by muscles located in the forearm, thereby broadening the biomechanical description of hand kinematics while remaining consistent with the HD sEMG acquisition framework. Specifically, the model will be extended to represent wrist supination and pronation, abduction and adduction, and flexion and extension, enabling a more comprehensive characterization of wrist biomechanics. In parallel, the dynamic neuromechanical interdependence among fingers during functional tasks will be systematically investigated using combined HD sEMG and kinematic data. Elucidating these interactions is expected to enhance the prediction of finger coordination patterns, support the development of more effective prosthetic and robotic control strategies, and provide deeper insight into the neural and biomechanical principles underlying skilled manual behavior.

## 5. CRediT Authorship Contribution Statement

**Valeria Avilés-Carrillo:** Data curation, Formal analysis, Investigation, Methodology, Software, Validation, Visualization, Writing – original draft. **Ricardo G. Molinari:** Conceptualization, Formal analysis, Methodology, Software, Validation, Visualization, Writing – original draft. **Guilherme A. G. De Villa:** Data curation, Investigation, Methodology, Software, Validation, Writing – review & editing. **Leonardo A. Elias:** Conceptualization, Funding acquisition, Methodology, Project administration, Resources, Supervision, Validation, Writing – review & editing.

## 6. Declaration of competing interest

The authors declare no known competing financial interests or personal relationships that could have influenced this work.

## 7. Acknowledgment

This work was supported by the Brazilian Public Ministry of Labor (MPT) under contract no. 002118.2019. CAPES supported Valeria Avilés-Carrillo under process no. 88887.842775/2023-00 (PROEX – Academic Excellence Program). Leonardo A. Elias is a Research Productivity Fellow of CNPq (process no. 316320/2023-4). Kinematic data were collected using the Vicon Motion Capture System, a multi-user system funded by FAPESP (proc. no. 2020/13293-0).

## Appendix A. RoM Post-hoc Comparisons

**Table A.1:** Bonferroni-corrected pairwise comparisons (*p*-values) between tasks for the thumb.

| Task | Index | Middle | Ring-little | Thumb | I. Pinch | M. Pinch | Tripod | 5-Finger |
| --- | --- | --- | --- | --- | --- | --- | --- | --- |
| Index |  | 0.573 | 0.163 | < **0.001** | < **0.001** | < **0.001** | < **0.001** | < **0.001** |
| Middle |  |  | 1.000 | < **0.001** | < **0.001** | < **0.001** | < **0.001** | < **0.001** |
| Ring-little |  |  |  | < **0.001** | < **0.001** | < **0.001** | < **0.001** | < **0.001** |
| Thumb |  |  |  |  | < **0.001** | < **0.001** | **0.004** | 1.000 |
| I. Pinch |  |  |  |  |  | 1.000 | 0.129 | < **0.001** |
| M. Pinch |  |  |  |  |  |  | 1.000 | **0.005** |
| Tripod |  |  |  |  |  |  |  | 0.182 |

Bold numbers indicate that $p < 0.05$ .

**Table A.2:**
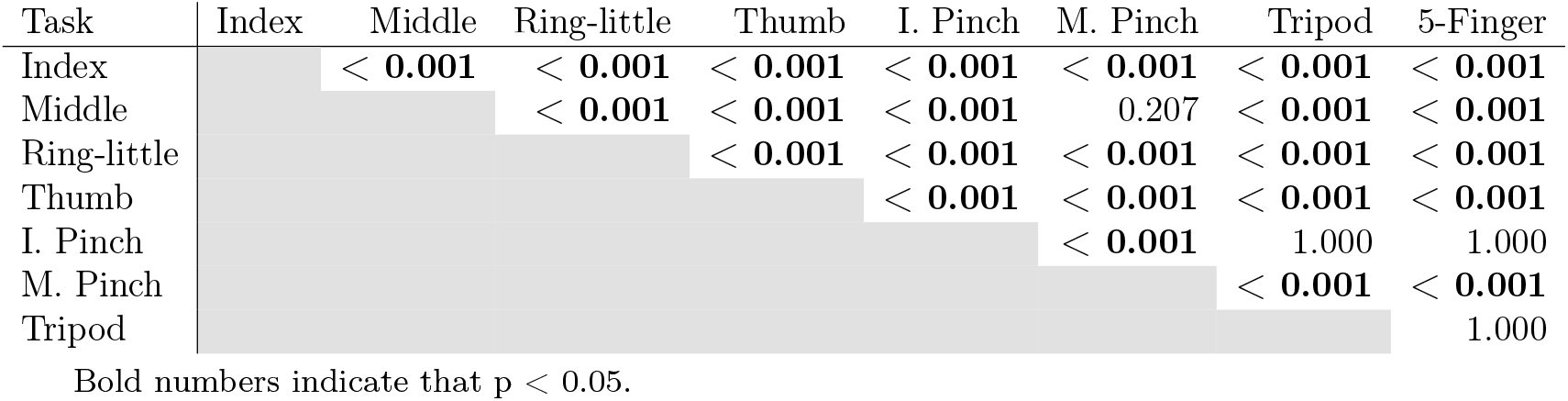
Bonferroni-corrected pairwise comparisons (*p*-values) between tasks for the index finger.

**Table A.3:** Bonferroni-corrected pairwise comparisons (*p*-values) between tasks for the middle finger.

| Task | Index | Middle | Ring-little | Thumb | I. Pinch | M. Pinch | Tripod | 5-Finger |
| --- | --- | --- | --- | --- | --- | --- | --- | --- |
| Index |  | < **0.001** | 0.074 | < **0.001** | **0.005** | < **0.001** | < **0.001** | < **0.001** |
| Middle |  |  | < **0.001** | < **0.001** | < **0.001** | < **0.001** | < **0.001** | < **0.001** |
| Ring-little |  |  |  | < **0.001** | < **0.001** | < **0.001** | < **0.001** | < **0.001** |
| Thumb |  |  |  |  | < **0.001** | < **0.001** | < **0.001** | < **0.001** |
| I. Pinch |  |  |  |  |  | < **0.001** | < **0.001** | < **0.001** |
| M. Pinch |  |  |  |  |  |  | 0.460 | 1.000 |
| Tripod |  |  |  |  |  |  |  | 0.409 |

Bold numbers indicate that $p < 0.05$ .

**Table A.4:**
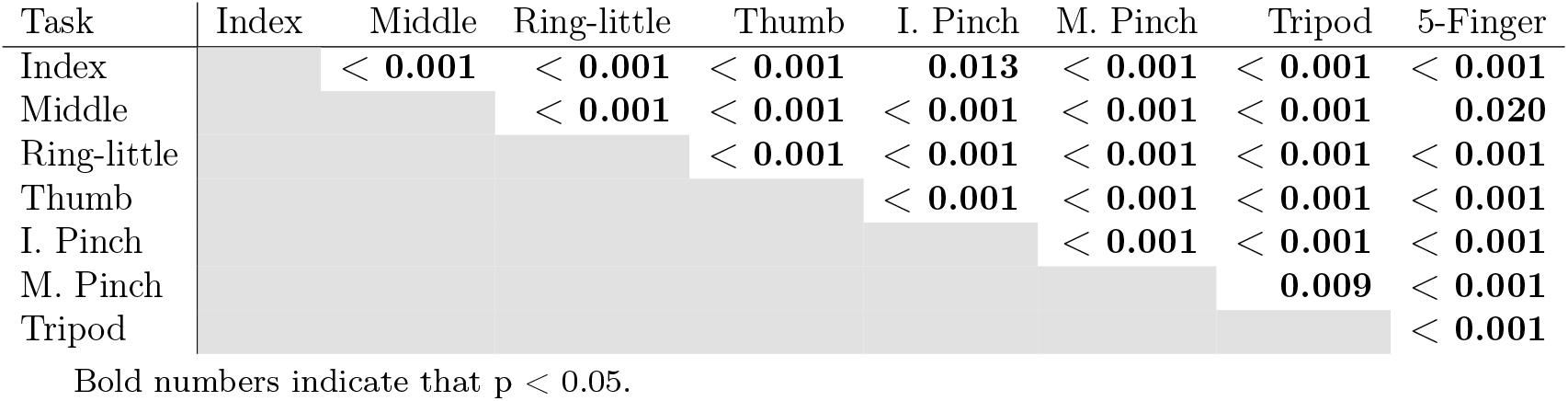
Bonferroni-corrected pairwise comparisons (*p*-values) between tasks for the ring finger.

| Task | Index | Middle | Ring-little | Thumb | I. Pinch | M. Pinch | Tripod | 5-Finger |
| --- | --- | --- | --- | --- | --- | --- | --- | --- |
| Index |  | < **0.001** | < **0.001** | < **0.001** | **0.013** | < **0.001** | < **0.001** | < **0.001** |
| Middle |  |  | < **0.001** | < **0.001** | < **0.001** | < **0.001** | < **0.001** | **0.020** |
| Ring-little |  |  |  | < **0.001** | < **0.001** | < **0.001** | < **0.001** | < **0.001** |
| Thumb |  |  |  |  | < **0.001** | < **0.001** | < **0.001** | < **0.001** |
| I. Pinch |  |  |  |  |  | < **0.001** | < **0.001** | < **0.001** |
| M. Pinch |  |  |  |  |  |  | **0.009** | < **0.001** |
| Tripod |  |  |  |  |  |  |  | < **0.001** |

Bold numbers indicate that $p < 0.05$ .

**Table A.5:** Bonferroni-corrected pairwise comparisons (*p*-values) between tasks for the little finger.

| Task | Index | Middle | Ring-little | Thumb | I. Pinch | M. Pinch | Tripod | 5-Finger |
| --- | --- | --- | --- | --- | --- | --- | --- | --- |
| Index |  | < **0.001** | < **0.001** | < **0.001** | 1.000 | < **0.001** | < **0.001** | < **0.001** |
| Middle |  |  | < **0.001** | < **0.001** | < **0.001** | 1.000 | 1.000 | < **0.001** |
| Ring-little |  |  |  | < **0.001** | < **0.001** | < **0.001** | < **0.001** | **0.002** |
| Thumb |  |  |  |  | < **0.001** | < **0.001** | < **0.001** | < **0.001** |
| I. Pinch |  |  |  |  |  | < **0.001** | < **0.001** | < **0.001** |
| M. Pinch |  |  |  |  |  |  | 1.000 | < **0.001** |
| Tripod |  |  |  |  |  |  |  | < **0.001** |

## Appendix B. Mean Speed Post-hoc Comparisons

**Table B.1:** Bonferroni-corrected pairwise comparisons (*p*-values) between 0.50 Hz and 0.75 Hz for all fingers within each Task (Freq*×*Task – Conditional on Task).

| Task | Thumb | Index | Middle | Ring | Little |
| --- | --- | --- | --- | --- | --- |
| Index | < **0.001** | < **0.001** | 0.055 | < **0.001** | < **0.001** |
| Middle | < **0.001** | **0.039** | < **0.001** | **0.008** | **0.016** |
| Ring-little | < **0.001** | < **0.001** | 0.760 | < **0.001** | < **0.001** |
| Thumb | < **0.001** | < **0.001** | < **0.001** | < **0.001** | < **0.001** |
| I. Pinch | **0.006** | < **0.001** | **0.002** | < **0.001** | < **0.001** |
| M. Pinch | **0.002** | **0.008** | 0.053 | 0.906 | **0.008** |
| Tripod | **0.002** | **0.001** | **0.024** | 0.448 | **0.006** |
| 5-Finger | < **0.001** | < **0.001** | **0.017** | < **0.001** | < **0.001** |

Bold numbers indicate that $p < 0.05$ .

## Appendix C. Normalized Total Harmonic Distortion post-hoc Comparisons

**Table C.1:** Bonferroni-corrected pairwise comparisons (*p*-values) between 0.50 Hz and 0.75 Hz for thumb, index, and ring fingers within each Task (Freq × Task – Conditional on Task).

| Task | Thumb | Index | Ring |
| --- | --- | --- | --- |
| Index | **0.035** | 0.314 | 0.299 |
| Middle | **0.002** | 0.261 | 0.296 |
| Ring-little | **0.001** | 0.057 | 0.072 |
| Thumb | 0.893 | **0.025** | **0.010** |
| I. Pinch | 0.494 | 0.454 | **0.031** |
| M. Pinch | 0.245 | 0.867 | 0.719 |
| Tripod | 0.470 | 0.160 | 0.683 |
| 5-Finger | 0.372 | 0.170 | 0.423 |

Bold numbers indicate that $p < 0.05$ .

**Table C.2:** Bonferroni-corrected pairwise comparisons (*p*-values) between tasks for the middle finger.

| Task | Index | Middle | Ring-little | Thumb | I. Pinch | M. Pinch | Tripod | 5-Finger |
| --- | --- | --- | --- | --- | --- | --- | --- | --- |
| Index |  | < **0.001** | **0.011** | < **0.001** | 1.000 | < **0.001** | < **0.001** | < **0.001** |
| Middle |  |  | < **0.001** | < **0.001** | < **0.001** | 1.000 | 1.000 | 1.000 |
| Ring-little |  |  |  | < **0.001** | 0.109 | < **0.001** | < **0.001** | < **0.001** |
| Thumb |  |  |  |  | < **0.001** | < **0.001** | < **0.001** | < **0.001** |
| I. Pinch |  |  |  |  |  | < **0.001** | < **0.001** | < **0.001** |
| M. Pinch |  |  |  |  |  |  | 1.000 | 1.000 |
| Tripod |  |  |  |  |  |  |  | 1.000 |

Bold numbers indicate that $p < 0.05$ .

**Table C.3:** Bonferroni-corrected pairwise comparisons (*p*-values) between tasks for the little finger.

| Task | Index | Middle | Ring-little | Thumb | I. Pinch | M. Pinch | Tripod | 5-Finger |
| --- | --- | --- | --- | --- | --- | --- | --- | --- |
| Index |  | < **0.001** | < **0.001** | 1.000 | 0.098 | < **0.001** | < **0.001** | < **0.001** |
| Middle |  |  | < **0.001** | < **0.001** | **0.032** | **0.026** | **0.001** | < **0.001** |
| Ring-little |  |  |  | < **0.001** | < **0.001** | < **0.001** | < **0.001** | 1.000 |
| Thumb |  |  |  |  | 0.230 | < **0.001** | < **0.001** | < **0.001** |
| I. Pinch |  |  |  |  |  | < **0.001** | < **0.001** | < **0.001** |
| M. Pinch |  |  |  |  |  |  | 1.000 | < **0.001** |
| Tripod |  |  |  |  |  |  |  | < **0.001** |

## Notes

https://doi.org/10.6084/m9.figshare.31032934

### Competing Interest Statement

The authors have declared no competing interest.

